# A high-impact *COL6A3* mutation alters the response of chondrocytes in neo-cartilage organoids to hyper-physiologic mechanical loading

**DOI:** 10.1101/2022.12.19.520461

**Authors:** Niek GC Bloks, Zainab Harissa, Shaunak S Adkar, Amanda R Dicks, Ghazaleh Hajmousa, Nancy Steward, Roman I. Koning, Aat Mulder, Berend B.R. de Koning, Margreet Kloppenburg, Rodrigo Coutinho de Almeida, Yolande FM Ramos, Farshid Guilak, Ingrid Meulenbelt

**Author notes:** Shared last author (Farshid Guilak, Ingrid Meulenbelt).

## Abstract

**Objectives:** The etiology of osteoarthritis revolves around the interplay between genetic predisposition and perturbing environmental cues, such as mechanical stress. The pericellular matrix, with its hallmark proteins collagen type VI and fibronectin, surrounds chondrocytes and is critical in transducing the biomechanical cues. The objective is to study the functional effects of an OA disease-risk mutation in *COL6A3* in interaction with hyper-physiological mechanical cues in a tailored human induced pluripotent stem cells (hiPSCs) derived cartilage organoid model.

**Method:** To identify pathogenic OA mutations exome sequencing in symptomatic OA patients was performed. To study functional effects, CRISPR-Cas9 genome engineering was used to introduce the mutation in our established human induced pluripotent stem cell-derived in-vitro neo-cartilage organoid model in interaction with hyper-physiological mechanical loading conditions.

**Results:** A high-impact mutation in *COL6A3* was identified that resulted in significantly lower binding between the PCM proteins COLVI and fibronectin (FN) and provoked an osteoarthritic chondrocyte state. Moreover, aberrant function of the PCM, secondary to the *COL6A3* mutation, abolished the initial stress responses marked particularly by upregulation of *PTGS2* encoding cyclooxygenase-2 (COX-2), after hyper-physiological mechanical loading conditions.

**Conclusion:** These findings demonstrate that ablating the characteristic transient COX-2 response after injurious mechanical cues may have a direct negative impact on chondrocyte health.

**What is already known:**

- The etiology of osteoarthritis revolves around the interplay between genetic predisposition and perturbing environmental cues, such as mechanical stress.
- The pericellular matrix, with its hallmark proteins collagen type VI and fibronectin, surrounds the chondrocytes and is critical in transducing biomechanical cues from the extracellular matrix to chondrocytes henceforth it determines the chondrocyte mechanical environment.
- The mechanical environment of the chondrocytes is a critical factor that influences chondrocyte health as it determines the balance between synthesis and degradation of the articular cartilage extracellular matrix.

**What this study adds:**

- A sustainable human induced pluripotent stem cell-derived in-vitro neo-cartilage organoid model that is tailored to study detailed biologic effects of mechanical cues to chondrocytes.
- An OA disease-risk mutation in *COL6A3* reduces the binding between collagen type VI to fibronectin and provoked an osteoarthritic chondrocyte state.
- Upon hyper-physiological mechanical loading, aberrant function of the pericellular matrix, secondary to the *COL6A3* mutation, ablates the initial transient inflammatory response, characterized particularly by *PTGS2* encoding cyclooxygenase-2 (COX-2).

**How this study might affect research practice or policy:**

- Inhibiting COX-2, as an important transient inflammatory response after hyper-physiological mechanical cues, could worsen the loss of structural integrity of the cartilage in osteoarthritis patients. Henceforth, prescription of COX-2 inhibitors as pain treatment for OA patients should be reconsidered.

## Introduction

Osteoarthritis (OA) is a complex, multifactorial disease revolving around the interplay between genetic and environmental risk factors. In particular, biomechanical cues play a critical role in not only joint health but also drive the onset and progression of disease [1]. Indeed, physiologic levels of mechanical loading by virtue of physical exercise can slow down OA disease progression [2, 3]. In contrast, hyper-physiological loading, as seen with post-traumatic injuries such as articular fracture, meniscal tear, or rupture of the anterior cruciate ligament, concomitant with altered joint kinematics, is a major risk factor for the onset and progression of OA [4]. Both physiological and hyper-physiological loading results in alterations in the structural composition of the cartilage extracellular matrix [1]. These findings suggest that the balance of chondrocyte anabolic and catabolic processes for the maintenance of cartilage is tightly regulated by biomechanical cues.

Chondrocytes reside in a pericellular matrix (PCM), which modulates the transduction of mechanical cues from the extracellular matrix (ECM) towards the chondrocyte. The PCM for that matter contains specific proteins such as collagen 6 (COLVI) and fibronectin (FN) [5]. In particular, collagen 6 (COLVI) is involved in regulating the mechanical properties of the PCM as well as calcium signaling in response to mechanical stress [5-7]. Knock-out of a COLVI sub-unit, *COL6A1,* in a murine model induces an osteoarthritic phenotype through dysregulation of mechano-transduction [6, 8]. While loss of COLVI in humans is characterized by joint contractures, hyperlaxity, and myopathy resulting in an inability to ambulate [9], the effect of aberrant COLVI function on chondrogenesis and mechano-transduction remains unclear.

Here we report the identification of a *COL6A3* missense mutation through exome sequencing of affected sibling pairs with generalized OA. *COL6A3,* coding for one of the monomeric sub-units of COLVI [10] that resides in [6] and interacts with other PCM proteins [11, 12], is likely involved in the regulation of cartilage structural composition and mechanical properties. We hypothesized that the identified *COL6A3* mutation would perturb cartilage mechano-transduction in response to hyper-physiologic loading, thereby contributing to OA pathogenesis. Recently, OA disease modeling platforms utilizing human induced pluripotent stem cells (hiPSCs) have been used to facilitate the interrogation of these pathogenic variants [13]. To gain an understanding of the role of COLVI in cartilage, we studied this mutation in an *in vitro* model of OA using hiPSC-derived cartilage. We employed genetically *COL6A3*-edited hiPSCs to an established *in vitro* cartilage organoid model, and these neo-cartilage organoids were exposed to hyper-physiological mechanical loading conditions. The effects of the mutation on COLVI’s interaction with other PCM proteins was studied. To study the downstream effects of this mutation in response to hyper-physiologic mechanical loading conditions, we characterized downstream molecular pathways based on transcriptome wide profiles. Understanding the role of COLVI in the regulation of mechanical stress will provide insight into etiology of OA and guide the development of effective treatment strategies for OA.

## Results

### Identification of a damaging mutation in COL6A3

Whole exome-sequencing was applied to a Caucasian sib pair of Dutch ancestry at age of 61 affected predominantly with symptomatic OA at multiple sites (GARP study) [14]. This resulted in the detection of 81,416 genetic variants, after which a prioritization schema was followed to identify pathogenic variants as previously described [15]. In short, first, we selected novel variants in in-house whole genome sequencing projects (N=222) and the BBMRI-Genome of the Netherlands project (GoNL, N=473) [15]. And second, damaging missense variants were selected based on the prediction of sorting intolerant from tolerant (SIFT). This prioritization generated a dataset of 38 novel coding variants that were predicted to have a functional impact on the protein **(table S1)**. And third, these 38 novel coding variants were then further prioritized based on their expression patterns in disease-relevant tissue by means of *in silico* analysis of 58 paired lesioned and preserved samples, resulting in two genes that are highly expressed in cartilage and differentially expressed between lesioned and preserved cartilage and bone; *MTHFR* (c.1667C>T, p.Pro597Leu; cartilage: FC=0.84, FDR=0.04, bone: not detected) and *COL6A3* (c.4510C>T, p.Arg1504Trp; cartilage: FC=1.75, FDR=2.25×10^-5^, bone: FC=1.69, FDR=0.042) [16]. *MTHFR* encodes for methylenetetrahydrofolate reductase, which is involved in folate metabolism [17]. *COL6A3* encodes for collagen VI subunit A3, which together with collagen VI subunit A1 and A2 forms a triple helical COLVI, a primary component of the cartilage PCM, and interacts with other proteins such as fibronectin and hyaluronan [11, 12, 18]. The identified COL6A3 mutation is located in the N-terminal VWA domains that are known to protrude away from the triple helical structure. The *COL6A3* mutation was selected based on COLVI’s role in regulating transduction of mechanical cues in cartilage towards chondrocytes [6]. To gain an understanding of aberrant COLVI function in cartilage, we selected this high-impact mutation for studying its effects in 3D in*-vitro* models of cartilage, and in interaction with hyper-physiological mechanical loading conditions.

### Introducing the mutation in hiPSCs

To investigate the effects of the aberrant COLVI on COLVI-mediated matrix deposition and molecular pathways, a gene-edited human induced pluripotent stem cell (hiPSC)-derived neo-cartilage organoid model was generated. We employed CRISPR-Cas9 single-stranded oligonucleotide-mediated homology-directed repair in hiPSCs to attain biallelic modification of rs144223596 (c.4510C>T) in an isogenic background, which was confirmed by Sanger sequencing **(Fig. S1)**. Although there was a small C peak visible at rs144223596, this was considered noise, and as such we considered the clone to be homozygous. Next, the *COL6A3*-edited and the unedited isogenic control hiPSCs were differentiated into chondrocytes using a previously established chondrogenic differentiation protocol [19]. In short, hiPSCs followed a step-wise differentiation protocol via mesodermal lineage differentiation towards chondroprogenitor cells [19]. These cells were then dissociated, and chondrogenesis was initiated in our organoid pellet model.

### Characterization of the matrix of isogenic control and COL6A3 mutant hiPSC-derived cartilage organoids

Successful differentiation towards chondrocytes and the production of neo-cartilage was confirmed by protein staining of collagen II (COLII), collagen VI (COLVI), and sulfated glycosaminoglycans (sGAGs) **(Fig. 1A)**. COLII and COLVI deposition was not affected by the mutation as measured by staining intensity **(Fig. 1B)**. However, quantification of sGAG deposition, using the dimethyl methylene blue (DMMB) assay normalized to DNA content, showed a significant reduction in the COLVI-mutant compared to the isogenic control neo-cartilage organoids **(Fig. 1C)**. Next, the isogenic and COLVI-mutant neo-cartilage organoids were characterized by targeted gene expression analysis using RT-qPCR of markers relevant for cartilage homeostasis and mechano-transduction **(table 1).** The *COL6A3* mutation reduced expression of the catabolic marker *ADAMTS5*, while also reducing expression of the anabolic markers *COL2A1* as well as *ACAN*.

**Table 1.** Effect of the COL6A3 mutation on markers of cartilage metabolism.

| Effect of *COL6A3* genotype |  |  |  |  |
| --- | --- | --- | --- | --- |
| Gene |  | beta | SE | *p*-value |
| *Catabolic* |  |  |  |  |
|  | *MMP13* | 0.02 | 0.27 | 0.94 |
|  | *MMP3* | -1.19 | 0.7 | 0.09 |
|  | *ADAMTS5* | -1.01 | 0.14 | <**0.001** |
| *Anabolic* |  |  |  |  |
|  | *COL2A1* | -0.47 | 0.2 | **0.02** |
|  | *ACAN* | -0.54 | 0.21 | **0.01** |
| *Hypertrophic* |  |  |  |  |
|  | *COL10A1* | -0.07 | 0.16 | 0.65 |

**Figure 1.**
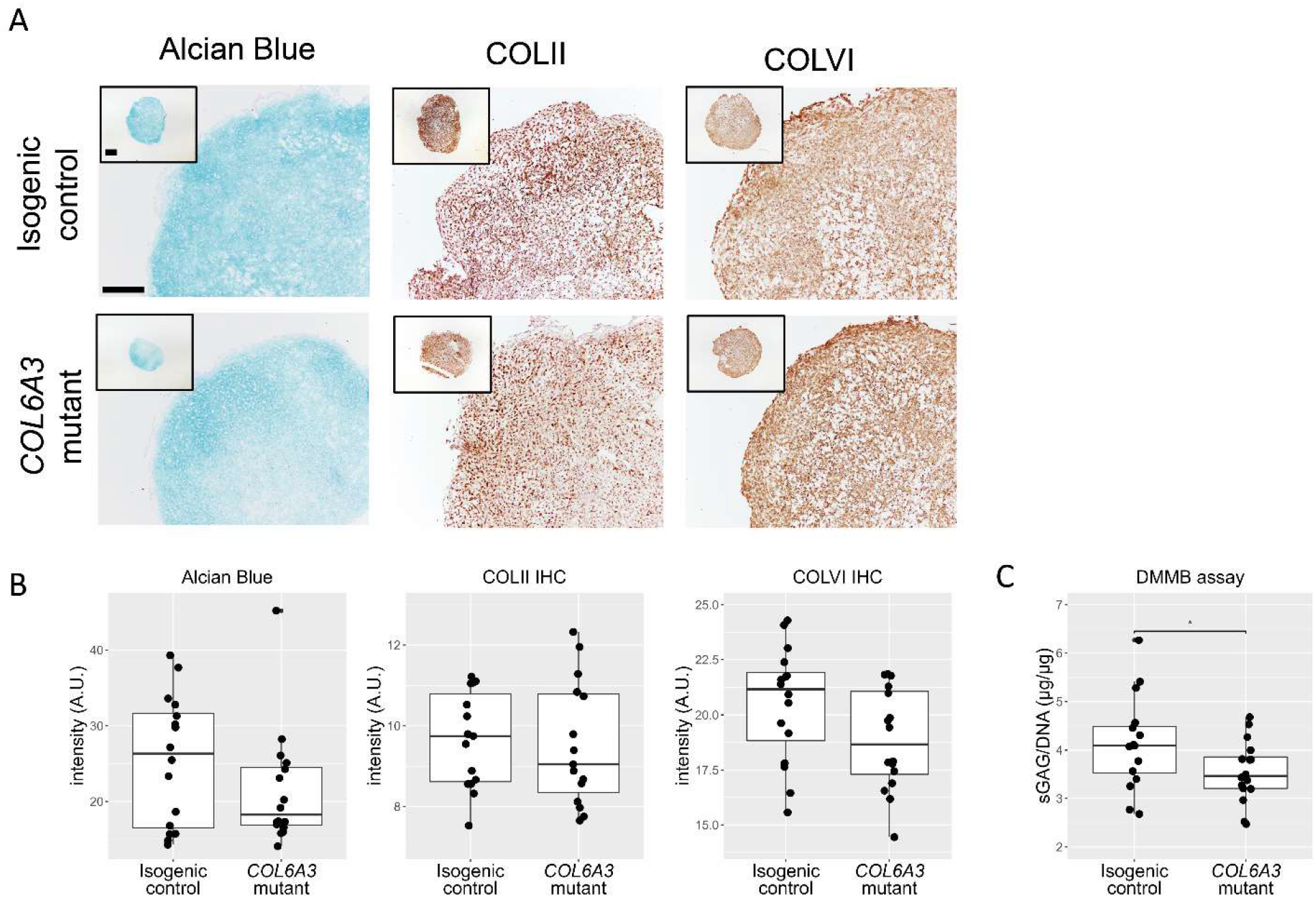
Effect of the mutation on neo-cartilage matrix deposition. **(A)** Representative images of Alcian blue staining marking sulfated glycosaminoglycans (sGAGs) and immunohistological staining of collagen II (COL II) and collagen VI (COLVI). Scale bar 200µm. **(B)** Quantification of Alcian blue, COLII, and COLVI in isogenic control and *COL6A3-*mutant neo-cartilage organoids showed no significant effect of the mutation (N=16). **(C)** Quantification of sGAG deposition in neo-cartilage organoids. sGAG deposition in COLVI*-*mutant neo-cartilage organoids is reduced in comparison to isogenic controls (beta=0.55 ± 0.28, P=4.9×10^-2^, N=16). Statistics are reported as beta ± standard error. The box plots represent 25th, 50th, and 75th percentiles, and whiskers extend to 1.5 times the interquartile range. Individual samples are depicted by black dots in each graph. P values were attained using a generalized linear model with intensity (for immuno-stainings) and (sGAGs/DNA) as dependent variable and genotype as independent variable.

### Transmission electron microscopy of the cartilage neo-matrix

To further investigate the effect of the mutation on the structural properties of the PCM and ECM, we performed transmission electron microscopy. As shown in **Figure 2A,** there was a reduced abundance of GAG-like structures in the *COL6A3*-mutant neo-cartilage organoids versus the isogenic control. Consistent with the DMMB assay, upon performing quantitative analysis of these GAG-like structures using a deep learning algorithm, we confirmed that mutant COLVI cartilage had significantly decreased aggregate size (Fig.2B)and GAG abundance (Fig. 2C).

**Figure 2.**
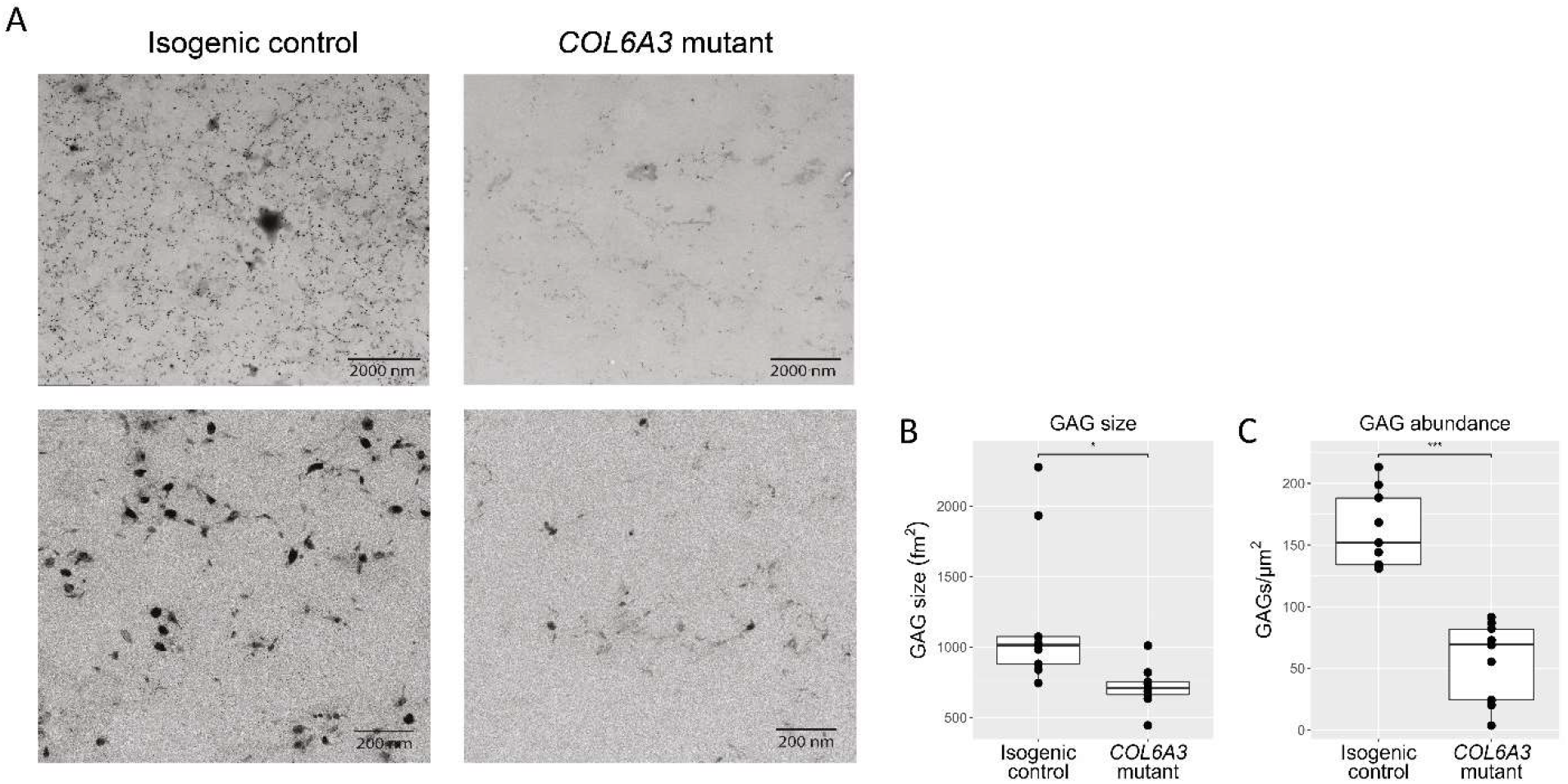
The COLVI mutation reduces sGAG aggregate size. **(A)** TEM image of isogenic control (left) and a *COL6A3*-mutant (right) neo-cartilage organoid. **(B)** Using a deep-learning algorithm, the sGAG aggregate size was measured, showed a reduction in the *COL6A3*-mutant neo-cartilage organoids (beta=-479.7, P=1.93X10^-2^). **(C)** GAG amount was decreased in the *COL6A3*-mutant neo-cartilage organoids (beta=-106.14 ± 14.95, P=2.52X10^-6^). Statistics are reported as beta ± standard error. The box plots represent 25th, 50th, and 75th percentiles, and whiskers extend to 1.5 times the interquartile range. Individual samples are depicted by black dots in each graph. P values were attained using a generalized linear model, with GAG size or GAGs/mm^2^ as a dependent variable and genotype as an independent variable.

### Binding of isogenic control and COL6A3 mutant COLVI to pericellular matrix proteins

The identified R1504W mutation is located in the N-terminal domain of COL6A3 coding for a von Willebrand Factor A domain, which is involved in binding two important proteins in cartilage PCM and ECM, fibronectin and hyaluronan [12, 13, 18]. Arginine at this position in the COL6A3 protein is evolutionary highly conserved, indicating that an amino acid change at this position is likely to affect protein function **(Fig. S2)**. Thus, we hypothesized that the change from a polar arginine to a non-polar tryptophan affects the interaction between COLVI and PCM/ECM proteins. Hereto, we have performed a fibronectin and hyaluronan solid-phase binding assay with wild-type and mutant COLVI, which was extracted from the chondrogenic differentiation medium of neo-cartilage organoids. Results, as shown in Figure 2, demonstrated that binding to fibronectin was reduced for mutant COLVI. However, we could not detect any binding of wild-type nor mutant COLVI to hyaluronan in our assay.

### Effect of missense COL6A3 mutation on the transcriptomic landscape

Next, using RNA sequencing, we determined the downstream transcriptional effects of these changes in the PCM and ECM related to the *COL6A3* mutation. The stranded RNA sequencing allowed us to confirm the genotyping as performed with Sanger sequencing **(Fig S1)**, which revealed the inclusion of both heterozygous (n=9) and homozygous (n=17) samples in our dataset To avoid compromising on power in addition to detecting the most robust differentially expressed genes (DEGs), we have pooled these heterozygous and homozygous samples after correction. Multifactorial analysis revealed 3700 significant DEGs between the *COL6A3* mutant and isogenic controls (FDR<0.05) **(Fig. 4A, table S2)**. Among these genes, 55% were upregulated and 45% were downregulated. Notable highly significant DEGs were related to development and cartilage metabolism, with the downregulation of structural PCM and ECM proteins, such as *COL27A1* and *PRG4*, increased catabolic activity, such as *MMP9*, and ECM mineralization, such as *SPP1* **(Fig. 4A-B)**.

**Figure 3.**
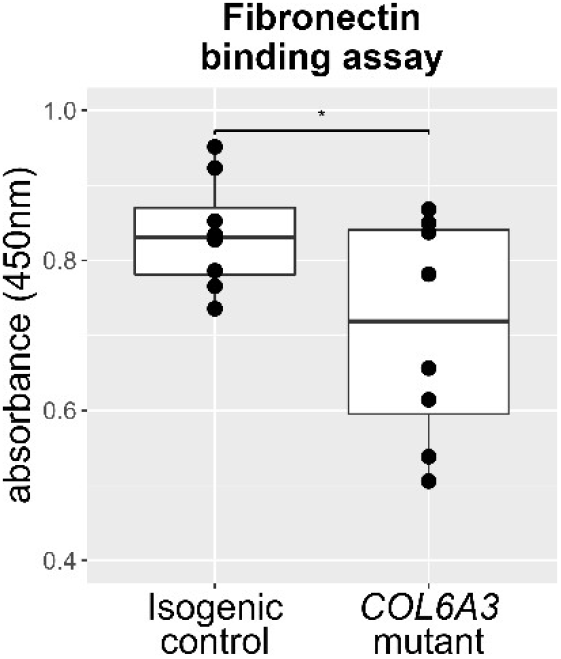
The COLVI mutation reduces binding to fibronectin. **(A)** Solid-phase binding assay with full-length fibronectin-coated wells and wild-type and R1504W mutant COLVI. R1504W showed a reduced binding to fibronectin (beta=-0.13 ± 0.06, P=4.37 x10^-2^, N=8). This assay did not show binding of COLVI to hyaluronan. Statistics are reported as beta ± standard error. The box plots represent 25th, 50th, and 75th percentiles, and whiskers extend to 1.5 times the interquartile range. Individual samples are depicted by black dots in each graph. P values were attained using a generalized linear model with absorbance at 450nm as dependent variable and genotype as independent variable.

**Figure 4.**
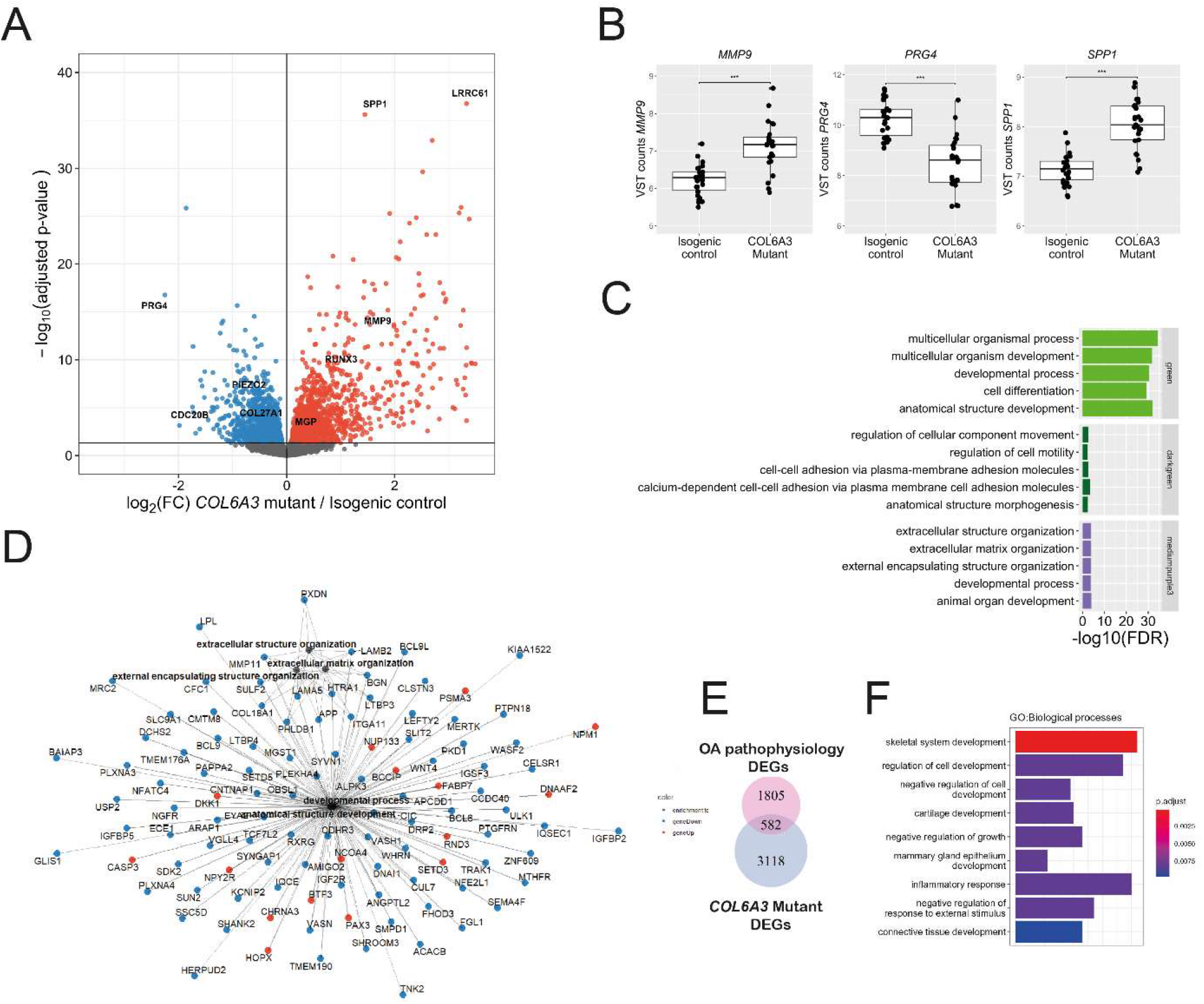
Transcriptomic profile in presence of the *COL6A3* mutation. **(A)** Volcano plot showing the differentially expressed genes (DEGs) in response to the *COL6A3* mutation (N=26). Red dots denote DEGs with an FDR<0.05 that are upregulated, and blue dots represent DEGs that are downregulated, as determined by DESeq2 analysis. **(B)** Notable examples of DEGs between isogenic control and *COL6A3*-mutant neo-cartilage pellets. The box plots represent 25th, 50th, and 75th percentiles, and whiskers extend to 1.5 times the interquartile range. Individual samples are depicted by black dots in each graph. ***FDR<0.001 **(C)** Overrepresentation enrichment analysis (KEGG, REACTOME, GO biological processes) of the top 3 weighted gene co-expression network analysis (WGCNA) co-expression modules where the first principal component is significantly associated with the *COL6A3* mutation. **(D)** Pathway – gene network of the enrichment analysis of the mediumpurple3 cluster. Lines depict the relationship between the genes and the pathways determined by enrichment analysis. Blue dots depict downregulated DEGs with the *COL6A3* mutation, while red dots depict upregulated DEGs with the *COL6A3* mutation. **(E)** Overlap between the *COL6A3* mutation DEGs and differential expression between lesioned and preserved cartilage from a previously published dataset (RAAK study, Ramos et al., 2014). **(F)** Over-representation enrichment analysis of overlapping DEGs between the mutation and lesioned versus preserved cartilage. Count depicts the number of genes that are categorized to each pathway.

To determine the biological processes associated with the *COL6A3* mutation, we performed a weighted gene co-expression network analysis (WGCNA) [20] on the RNA-sequencing data set. This resulted in the detection of 20 distinct co-expression networks **(Fig. S3)**. Multifactorial regression analysis revealed 10 co-expression networks with the first principal component significantly associated with the *COL6A3* genotype **(Fig. S3)**. The top 3 most significant co-expression networks associated with the *COL6A3*-mutant genotype were enriched for developmental processes, cell-cell adhesion/anatomical structure morphogenesis, and ECM organization **(Fig. 4C)**. Enrichment of ECM organization again underlines the detrimental effects of the *COL6A3* mutation on the development of cartilage, affecting genes such as *LAMA5*, *LAMB2,* and *COL18A1* that code for structural PCM components **(Fig. 4D)**. While enrichment of cell-cell adhesion pathways suggests that the mutation also affects mechano-transduction via altered cell-cell adhesion by downregulation of cadherin-associated genes, such as *CDH18*, *CDH6*, *PTGER4*, *MAPK14,* and *ADAMTS5* **(Fig. S4)**. Of note is that *ADAMTS5* is downregulated in the *COL6A3* mutant while quantification of sGAGs, the target peptides of ADAMTS5 cleavage, are reduced at both the gene level (*ACAN*, **Table 1**) and protein level **(Fig. 1C; Fig. 3A-C).**

To study the relevance of the *COL6A3* mutation to OA pathophysiology, we determined the overlap of gene expression between the effects of the *COL6A3* mutation and the differential expression between lesioned and preserved cartilage from patient material collected post-joint replacement surgery in a previously published dataset containing 58 paired samples [16] revealing a subset of 582 overlapping DEGs **(table S4)**. We then performed pathway analysis on this subset of genes **(Fig. 4B)**. The most significant enriched pathway was skeletal system development, including transcription factors (*RUNX3*, *TNFRSF11B*), ECM components (such as *COL27A1*, *COL11A2*), and anabolic factors (such as *IGF1*, *BMP3*). Well-known pathways related to inflammation “inflammatory response” and “acute inflammatory response” were also enriched containing inflammatory cytokines (IL31a) and general inflammatory response factors (*PTGES*, *HLA-e*). Also, the “ossification” pathway was enriched, containing OA risk genes (*SPP1*, *TNC,* and *MGP*). Genes related to cartilage metabolism were confirmed by quantitative reverse transcription polymerase chain reaction (RT-qPCR) **(table S5).** Together, this data suggests that the *COL6A3* mutation results in downstream expression changes, in part describing the osteoarthritis pathophysiology.

Additionally, to explore the effect of the hetero- and homozygous *COL6A3*-mutated hiPSC-derived neo-cartilage organoids, a dose-response effect was modeled for DEGs (FDR<0.05) responding to the *COL6A3* mutation. To this end, the hetero- and homozygous *COL6A3* mutation was considered as a covariate and analyzed for differential expression. This revealed a robust dose-response effect (FDR<0.05) of the *COL6A3* mutation on gene expression in 2610 genes out of the 3700 DEGs affected by the mutation, **(table S6; Figure S6**), such as MMP9, PRG4, and SPP1 This dose-response effect further underlines the biological validity of the gene expression changes due to *COL6A3* mutation.

Taken together, we showed that the *COL6A3* mutation reduces sGAG deposition **(Fig. 1)**, reduces binding to fibronectin **(Fig. 2)**, and lowers the expression of mechano-sensor genes **(table 1)** suggesting the mutation could affect the transduction of mechanical cues. Hence, we determined whether the *COL6A3* mutation affects the transcriptomic response to hyper-physiologic mechanical loading conditions. Hereto, two different organoid models were applied; cylindrical constructs in which hiPSC-derived pellets were digested to obtain single-cell chondrocytes that were then encapsulated in an agarose gel, and spherical constructs with neo-cartilage deposited by hiPSC-derived chondrocytes. Both these organoid models were exposed to hyper-physiologic mechanical loading conditions [20% sinusoidal peak-to-peak strain at 5hz for 10 minutes]. Transcriptional activity was measured 12 hours after mechanical stimulation in isogenic controls (unloaded isogenic controls - n=13, mechanically stimulated isogenic controls - n=13) as well as *COL6A3*-mutant (unloaded *COL6A3* mutants – n=14 and mechanically loaded *COL6A3* mutants - n=12) organoids.

### Effect of mechanical loading on the transcriptomic landscape

Next, we characterized the effect of hyper-physiologic loading conditions [21] on these neo-cartilage organoids. Multifactorial analyses revealed 177 DEGs (FDR<0.05) between unloaded and mechanically loaded organoids, of which 74% were upregulated and 26% were downregulated **(Fig.6A, table S7)**. Notable genes were, amongst others; *CD44*, *CAV1*, *ITGA5* **Fig. 6B**), which are all genes encoding for mechano-sensors involved in OA pathophysiology.

**Figure 5.**
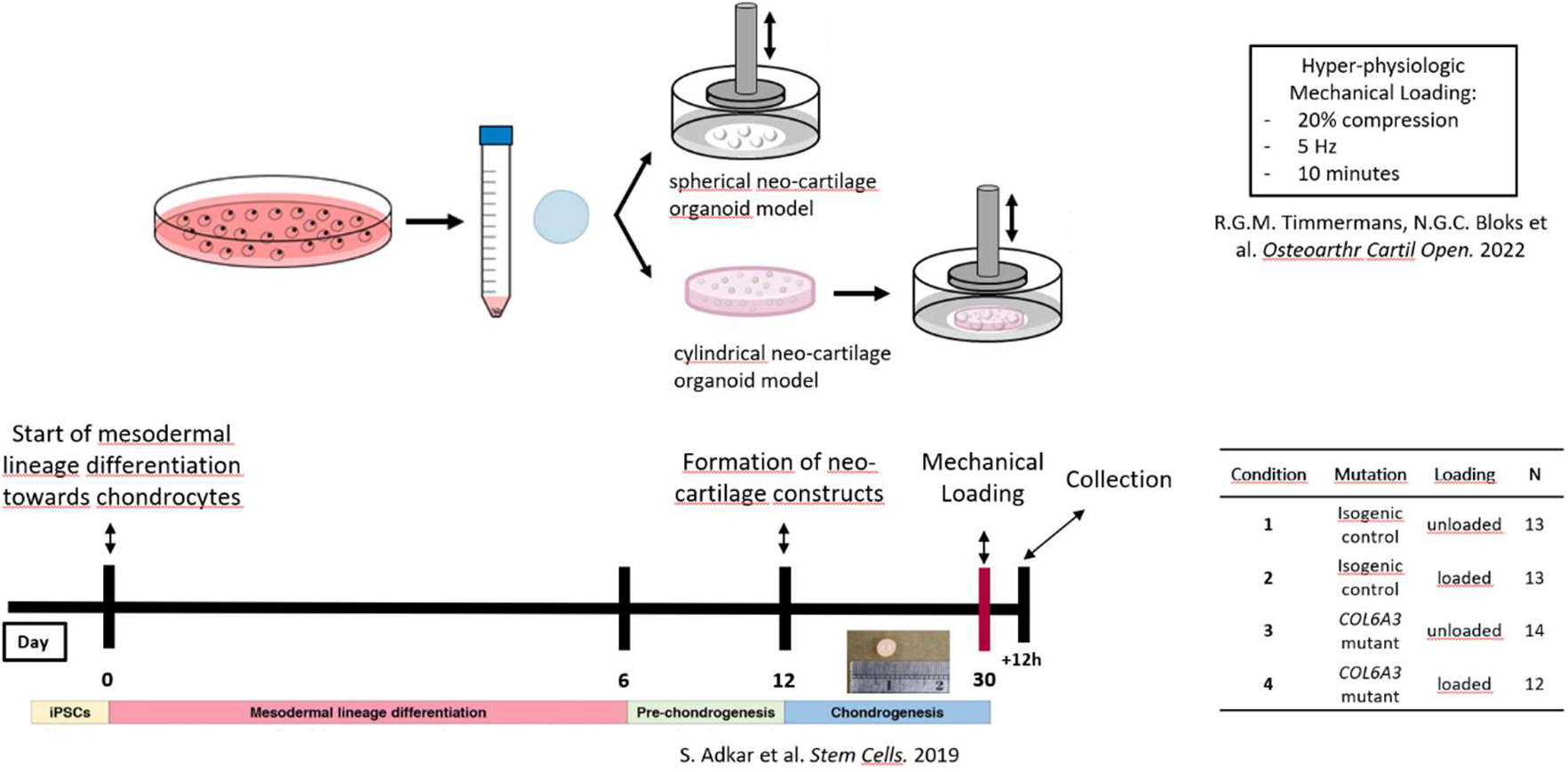
Chondrogenic models to study the effects of aberrant COLVI function in interaction with hyper-physiologic mechanical loading conditions. hiPSCs in which the R1504W mutation was introduced using CRISPR-Cas9 genome editing. These cells were differentiated using an established differentiation protocol to produce neo-cartilage organoids. Two different organoid models were employed; 1. A spherical pellet model harnessing the original matrix produced by the hiPSCs. 2. A cylindrical organoid model in which the hiPSc-derived chondrocytes were embedded in an agarose construct, ideally suited for testing the effects of mechanical loading conditions. These constructs were both exposed to hyper-physiological loading conditions, after which the organoids were harvested for downstream analysis.

**Figure 6.**
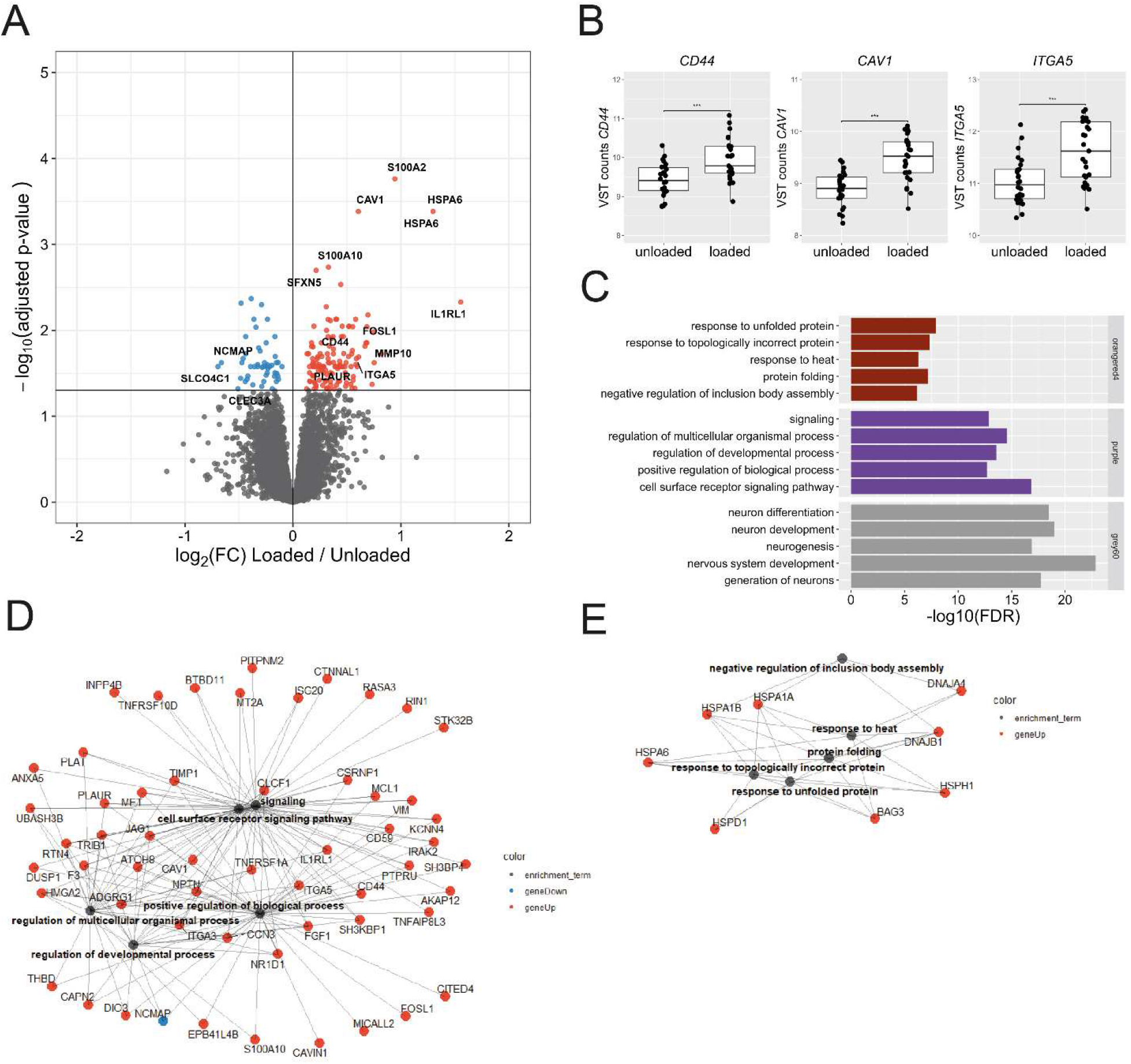
Transcriptomic profile in response to hyper-physiologic mechanical loading conditions. **(A)** Volcano plot of differentially expressed genes (DEGs) in response to hyper-physiologic loading conditions. Red dots denote differentially DEGs with an FDR<0.05 that are upregulated, and blue dots represent DEGs that are downregulated as determined by DESeq2 analysis. **(B)** Notable examples of mechano-sensor genes upregulated in response to hyper-physiological mechanical loading conditions. The box plots represent the 25th, 50th, and 75th percentiles, and the whiskers extend to 1.5 times the interquartile range. Individual samples are depicted by black dots in each graph. *FDR<0.05 **(C)** Overrepresentation enrichment analysis (KEGG, REACTOME, GO biological processes) of the top 3 weighted gene co-expression network analysis (WGCNA) co-expression modules of which the first principle component is significantly associated with hyper-physiologic mechanical loading conditions. **(D-E)** Gene-pathway network of the enrichment analysis of the purple **(D)** and the orangered4 **(E)** co-expression network where lines depict the relationship between the genes and the pathways determined by enrichment analysis. Blue dots depict downregulated DEGs in response to hyper-physiologic mechanical loading conditions, red dots depict upregulated DEGs in response to hyper-physiologic mechanical loading conditions.

To determine the biological processes affected by hyper-physiologic mechanical loading WGCNA co-expression networks were associated with mechanical loading, revealing 5 significantly associated co-expression networks **(Fig. S2; table S3)**. The top 3 most significantly associated modules were enriched for developmental and signaling processes, stress responses, and neuronal pathways **(Fig. 6B)**. While enrichment of signaling responses containing genes such as *IL1RL1* and *FOSL1* show an adaptive response to hyper-physiologic mechanical loading conditions **(Fig. 6D)**, enrichment of stress responses with genes such as *HSPA1B* and *DNAJA4* underlines the damaging effects of the hyper-physiologic mechanical loading conditions **(Fig 6E).**

### Effect of the COL6A3 missense mutation on the response to hyper-physiological mechanical loading conditions

Finally, the effects of the *COL6A3* mutation on the response to hyper-physiological mechanical loading were investigated. To this end, a multifactorial analysis was performed resulting in a set of 135 genes with a significant interaction effect (P<0.01) indicating that the mutation affected the response to hyper-physiological mechanical loading **(table S8)**. Of these 135 genes, 70 proteins show a significant protein-protein interaction (PPI) (FDR<0.05) as determined by STRING-DB **(Fig. 7A)**. Notable is that highly connected genes in this PPI network such as; *PTGS2*, *IL1R1, IRAK2, PECAM1,* and *ADAMTS5* are all related to catabolic and inflammatory signaling. *PTGS2* is known to be regulated by TRPV4 signaling in response to mechanical stress [22] and is involved in inducing an inflammatory response to stimuli [23]. *IL1R1* codes for the receptor of interleukin-1, one of the key inflammatory markers in OA, and thus is involved in inflammatory signaling, while *IRAK2* encodes for the interleukin-1 receptor-associated kinase 2, which is involved in interleukin-1 (IL-1) induced upregulation of NF-κβ [24]. Upon performing stratified analysis, it was shown that the interaction effect was particularly caused by an upregulation of gene expression observed in isogenic controls in response to hyper-physiological loading conditions, that appeared absent in the *COL6A3* mutant neo-cartilage organoids **(Figure 7B; table S9).**

**Figure 7.**
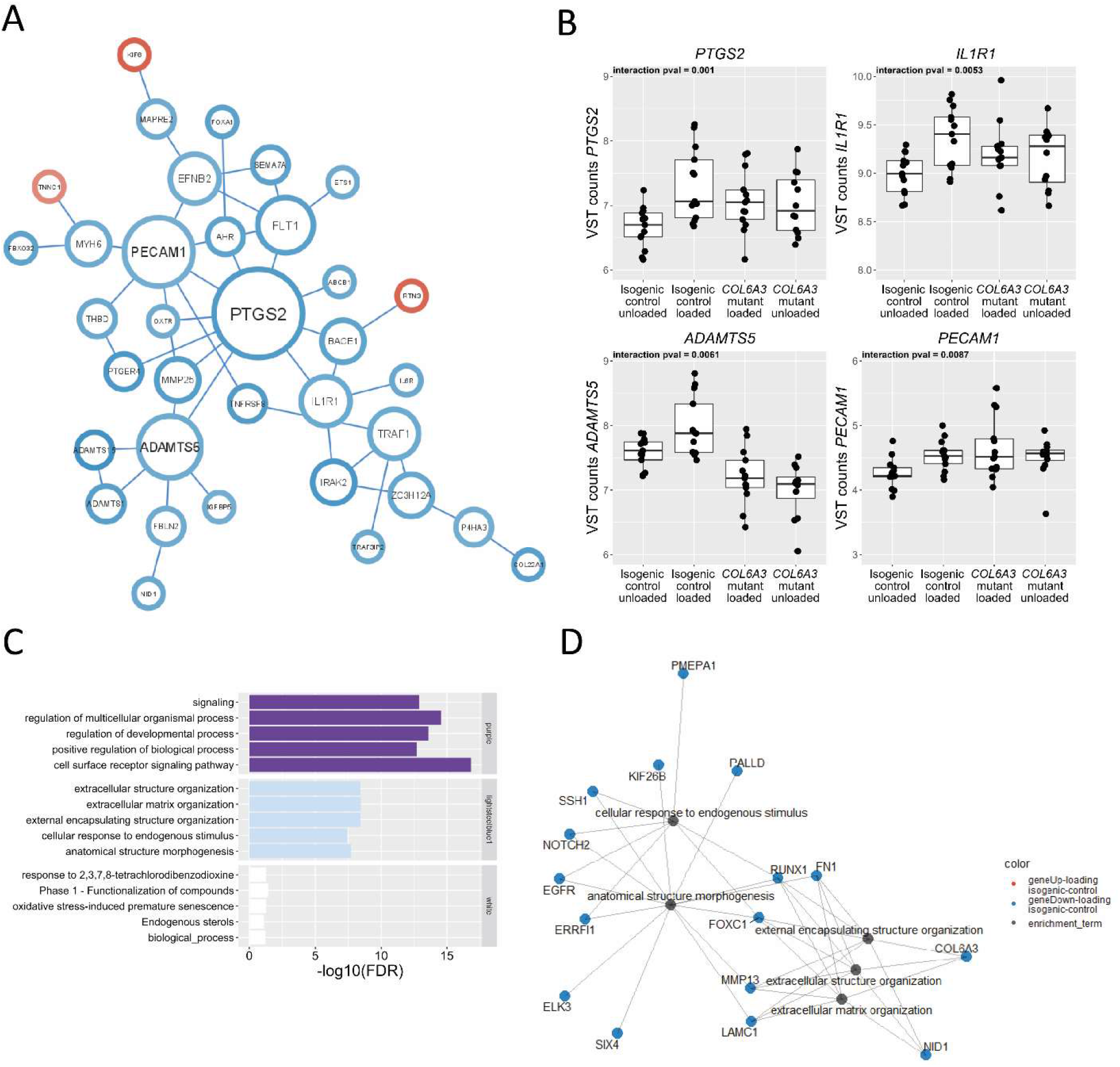
The *COL6A3* mutation affects the biological response to hyper-physiologic loading conditions. **(A)** Protein-protein interaction (PPI) network based on the STRING-DB of genes that show an interaction (P*<*0.01) between the *COL6A3* mutation and hyper-physiological mechanical loading conditions. Red circles denote an increased response to hyper-physiological mechanical loading due to the *COL6A3* mutation, while blue circles denote a reduced response to mechanical stress due to the *COL6A3* mutation (only showing connected nodes). Relative node size depicts number of connections for each gene within the network **(B)** Examples of central DEGs in the PPI that are related to an inflammatory response. The box plots represent 25th, 50th, and 75th percentiles, and whiskers extend to 1.5 times the interquartile range. Individual samples are depicted by black dots in each graph. **(C)** Gene enrichment analysis of WGCNA hubs significantly related to the interaction between the *COL6A3* mutation and the response to hyper-physiological mechanical loading conditions. **(D)** Pathway – gene network of the enrichment analysis of the purple cluster. Lines depict the relationship between the genes and the pathways determined by enrichment analysis. Blue and red dots depict, respectively, downregulated and upregulated DEGs in response to hyper-physiological loading conditions in the isogenic control neo-cartilage organoids.

To study the biological processes in which the response to hyper-physiological loading differed between the isogenic control and the *COL6A3* mutants, associate this interaction effect with the principal component of each detected co-expression network. This revealed three distinct co-expression networks associated with the interaction effect, which were enriched for processes related to signaling, ECM organization, and general biological processes **(Figure 7C)**. Enrichment of pathways related to signaling shows an aberrant response in genes such as *INHBA* which acts downstream of TRPV4 calcium channels [22], *CAV1*, *SEMA7A*, and *MMP3,* which are all genes that respond to mechanical loading in isogenic controls, with an absent loading effect in the *COL6A3* mutant **(table S3)**. Again, the enrichment of processes related to ECM formation and organization, containing genes such as; *MMP13, RUNX1, FN1, LAMC1, EGFR,* and *PMEPA1* **(Fig. 7D)**, underlines the effect of an impaired repair response to hyper-physiological mechanical loading conditions in the *COL6A3* mutant. Also, the enrichment of general biological processes including *IL1R1* highlights the aberrant response to mechanical loading in the *COL6A3* mutant neo-cartilage organoids.

## Discussion

In the current study, we identified a missense mutation in *COL6A3* (c.4510C>T, p.Arg1504Trp) in a sibling pair of the GARP-study affected with symptomatic OA in two or more joint sites [14]. By introducing this high-impact mutation in hiPSCs using CRISPR-Cas9 genome engineering while employing these cells in an established 3D *in vitro* neo-cartilage organoid model, we showed that the mutation decreases cartilage matrix integrity, as reflected by a reduction in abundance and size of GAGs. Moreover, by subsequently isolating mutated COL6 protein, we showed that the mutation reduced binding to fibronectin. Analysis of the transcriptome-wide gene expression changes with *COL6A3* mutation showed overlap with transcriptomic changes due to OA-pathophysiology [16] indicating that an osteoarthritic chondrocyte state appeared secondary to alterations in PCM function due to mutant *COL6A3.* Finally, by exposing the neo-cartilage organoids to hyper-physiological mechanical stress, we demonstrated that mutated *COL6A3* abolished the characteristic upregulation of inflammatory signaling after mechanical loading [21, 22, 25] with *PTGS2*, *PECAM1*, and *ADAMTS5*, as most central and notable genes. Taken together, our findings suggest that the identified mutation in COL6A3 resulted in impaired binding between COLVI and the PCM protein fibronectin and abolished the initial stress response to hyper-physiologic mechanical loading conditions that could affect the propensity of the chondrocyte to enter an OA disease state.

Although it has been suggested that *PTGS2*, commonly known as cyclooxygenase-2 (COX-2), as part of the PGE2–EP4 axis, is essential to initiate repair in other tissues in response to stressors [26], we here showed for the first time that absence of this COX-2 response could underlie OA pathophysiology. Notable in this respect is that COX-2 inhibitors are frequently prescribed for OA pain [27]. In light of our result, we advocate that increased joint loading upon prescription of COX-2 inhibitors for OA pain could worsen OA pathophysiology.

By introducing an high-impact *COL6A3* mutation in a hiPSC-derived neo-cartilage organoid model that had been genome-edited via CRISPR-Cas9 to harbor this mutation, we have been able to study in detail chondrocyte mechano-biologic effects to hyper-physiological mechanical stress. Nonetheless it should be noted that upon checking the homozygous introduction of the *COL6A3* mutation by genotyping the RNA sequencing data, heterozygous *COL6A3* mutated samples were identified. These heterozygous *COL6A3* mutant cells could have arisen due to a selective advantage upon expanding hiPSCs. Availability of hetero- and homozygous samples, however, did allow us to study a possible dose-response effect of the mutation on RNAseq data and confirmed a dose-response effect in 2610 genes out of the 3700 DEGs. This adds to the validity of our data and biological relevance of the studied *COL6A3* mutation, as there was an additive effect of the hetero- and homozygous *COL6A3* mutants on DEGs such as SPP1, MMP9, and PGR4 **(table S6, Fig. S7)**.

We have applied both cylindrical disc-shaped organoids [22] as well as spherical pellets [13, 21]. The advantage of the first model is an ensured equal distribution of mechanical stress throughout the disc-shaped sample. The advantage of the second model is that it allows harnessing the original matrix deposited by the hiPSC-derived chondrocytes thereby permitting the characterization of changes in the ECM. Additionally, this model of hyper-physiological loading in hiPSC neo-cartilage organoids can be expanded to define and apply more beneficial mechanical loading regimes, further unraveling the shift in molecular mechanisms underlying the normal physiological response to loading that might further aid in the development of treatment strategies.

In contrast to previous findings [12], we could not detect any binding of COLVI with hyaluronan, which might be explained by the source of the COLVI proteins, as they were extracted from the culture medium. Alternatively, other intermediary proteins are necessary for binding COLVI to hyaluronan which might not have been present in the medium [11]. Moreover, we have used TEM in combination with a, yet to be published, machine-learning tool to detect changes in GAG size and abundance. We relied on the similarity with previously published data to train the machine-learning algorithm. Nevertheless, the reduced abundance of GAGs as measured with TEM is in line with the results of the DMMB assay, as well as the Alcian blue staining.

By using genetic engineering of hiPSC-derived neo-cartilage organoids while implementing hyper-physiological mechanical loading conditions, we established a tailored model to study the biological function of proteins in the transduction of mechanical cues from ECM to chondrocytes. By using this model, we here showed that an identified OA disease-risk mutation in COL6A3 reduces the binding between collagen type VI to fibronectin and provoked an osteoarthritic chondrocyte state. Moreover, we are among the first to demonstrate that aberrant function of COLVI in the PCM abolishes the initial inflammatory response to hyper-physiological loading, as marked by, amongst others, changes in *PTGS2* encoding COX-2. In light of these finding, we advocate that COX-2 inhibitors prescribed as pain medication to OA patients could have a negative impact on chondrocyte health when patients are exposed to hyper-physiological mechanical loading.

## Materials and Methods

### Exome sequencing

Exome sequencing of a sibling pair with generalized OA at multiple joint sites was performed by Illumina HiSeq 2000 technology (Beijing Genome Institute) using the protocol described in the supplementary methods.

### hiPSC line and cell culture

An hiPSC line as described earlier was used as the unedited isogenic control [19]. In short, RVR-iPSC line was retrovirally reprogrammed from BJ fibroblasts and characterized. Further culture conditions can be found in the supplementary methods.

### Genome editing of hiPSCs

We employed CRISPR-Cas9 single-stranded oligonucleotide-mediated homology-directed repair in hiPSCs to attain biallelic modification of rs144223596 (c.4510C>T) in an isogenic background, Additional information on the genomic target, guide sequences and transfection protocol can be found in the supplementary methods.

### hiPSC differentiation to induced chondrocytes

Generation of induced chondroprogenitor cells (hiCPCs) was based on a protocol previously described [19] which was shown to produce similar neo-cartilage to that produced by human primary articular chondrocytes [28] further described in the supplementary methods.

### Mechanical loading

The spherical-shaped neo-cartilage constructs were mechanically loaded using a MACH-1 mechanical testing device (Biomomentum), at a rate of 5hz with 20% sinusoidal peak-to-peak strain for 10 minutes as described earlier [21]

### sGAG measurement

Sulfated glycosaminoglycan (sGAG) concentrations in the neo-cartilage organoids (µg sGAG/µg DNA) were measured using the Farndale Dimethyl Methylene Blue (DMMB, Sigma) method [29]. Additional information can be found in the supplementary methods.

### Histology and immunohistochemistry

Neo-cartilage samples were fixed in 4% formaldehyde and embedded in paraffin. Sections were stained with Alcian Blue (Sigma-Aldrich) and Nuclear Fast Red (Sigma-Aldrich). Deposition of collagen II and collagen VI in the neo-cartilage constructs was visualized immunohistochemically according to the protocol described in the supplementary methods.

### RT-qPCR

Per sample, two replicate mRNA samples were measured in triplicates in a MicroAmp™ Optical 384-Well Reaction Plate (ThermoFisher Scientific), using the QuantStudio™ Flex Real-Time PCR system (Applied Biosystems™). Additional information can be found in the supplementary methods.

### RNAseq

RNA from neo-cartilage constructs was extracted 12 hours post mechanical loading and analyzed using the Illumina NOVAseq 6000. Additional information on RNA isolation, mapping, alignment, and data processing and analysis can be found in the supplementary methods.

### Solid-phase binding assay

Conditioned medium of wild-type and *COL6A3* mutant organoids was collected and concentrated in preparation for the binding assay. To this end, 450 μl of medium was collected in 100 K molecular weight cutoff Pierce Protein Concentrators (Thermo Scientific) and centrifuged for 10 min at 12,000g. Subsequently, COL6 concentration was determined using the Human COL6A3 ELISA Kit (Assay Genie) according to the manufacturer’s protocol which can be found in the supplementary methods.

### Transmission electron microscopy

To the neo-cartilage organoids, consisting of cells and matrix, a previously established protocol was used to perform TEM. Additional information on sample processing, image acquisition, processing and analysis can be found in the supplementary methods.

### Statistical analysis

For all data analysis except RNA sequencing we have used a generalized linear model including the factors hyper-physiological loading and the *COL6A3* mutation using R statistical software version 4.1.1.

## Author Contributions

N.G.C.B., Z.H., A.R.D., Y.F.M.H, F.G., I.M. developed the concept of this study; N.G.C.B., Z.H., S.S.A., A.R.D., G.H., N.S. M.K., R.I.K., A.A.M., B.B.R.K. and F.G., acquired materials and data; N.G.C.B., R.C.D.A., B.B.R.K., Y.F.M.H, F.G. and I.M. analyzed the data; and all authors contributed to the writing of the manuscript.

## Data and materials availability

All data needed to evaluate the conclusions in the paper are present in the paper and/or the Supplementary Materials.

