## Supplementary Materials for "A high-impact *COL6A3* mutation alters the response of chondrocytes in neo-cartilage organoids to hyper-physiologic mechanical loading"

Niek G.C. Bloks et al.

Corresponding author: Ingrid Meulenbelt,


**This PDF file includes:**

Figs. S1 to S7

**Other Supplementary Materials for this manuscript include the following:**

Tables S1-S10

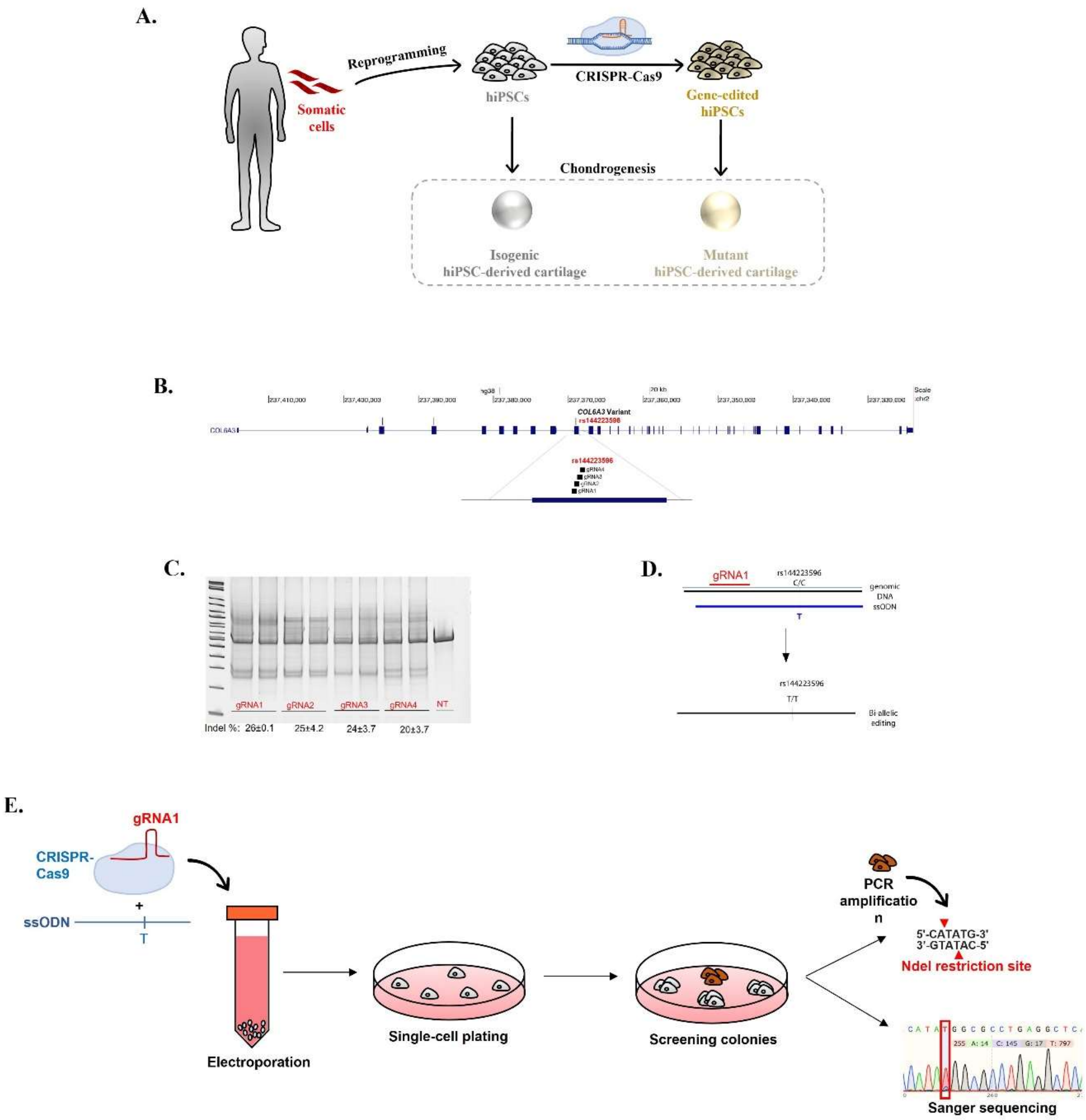


**Supplementary Figure 1.** Overview of experimental set-up CRISPR-Cas9 gene editing of rs144223596 C>T. **(A)** Schematic of hiPSCs editing OA variants. hiPSCs are gene-edited using CRIPSR/Cas9 to introduce OA disease variants found using exome sequencing studies. Gene-edited hiPSCs alongside their isogenic controls are then differentiated into chondrogenic lineage to obtain hiPSC-derived cartilage, which can be used in downstream analysis to determine the mechanisms linking the mutation to OA. **(B)**. gRNAs were designed targeting genomic DNA flanking the COl6A3 variant rs144223596 (http://genome.ucsc.edu) *(34)* and **(C)**. Screened in HEK293T cells to evaluate cutting efficiency using Surveyor assay. **(D)**. Schematic of biallelic editing using optimized gRNA targeting the risk variant and ssODN harboring T risk allele. **(E)** Schematic of single-cell expansion into colonies following electroporation with gRNA complexed to Cas9 and ssODN. Colonies were screened by PCR amplification and digestion with an NdeI and evaluated using sanger sequencing

A

| substitution | preservation time | Message | Pdel |
|---|---|---|---|
| R1504W | 361 | possibly damaging | 0.5 |

B

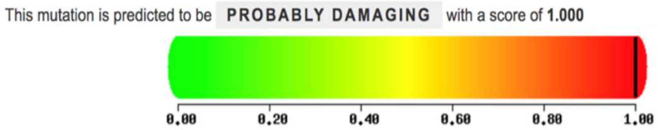


**Supplementary Figure 2**. In-situ predictions of damaging effects COL6A3 R1504W mutation **(A)** Panther prediction **(B)** Polyphen 2

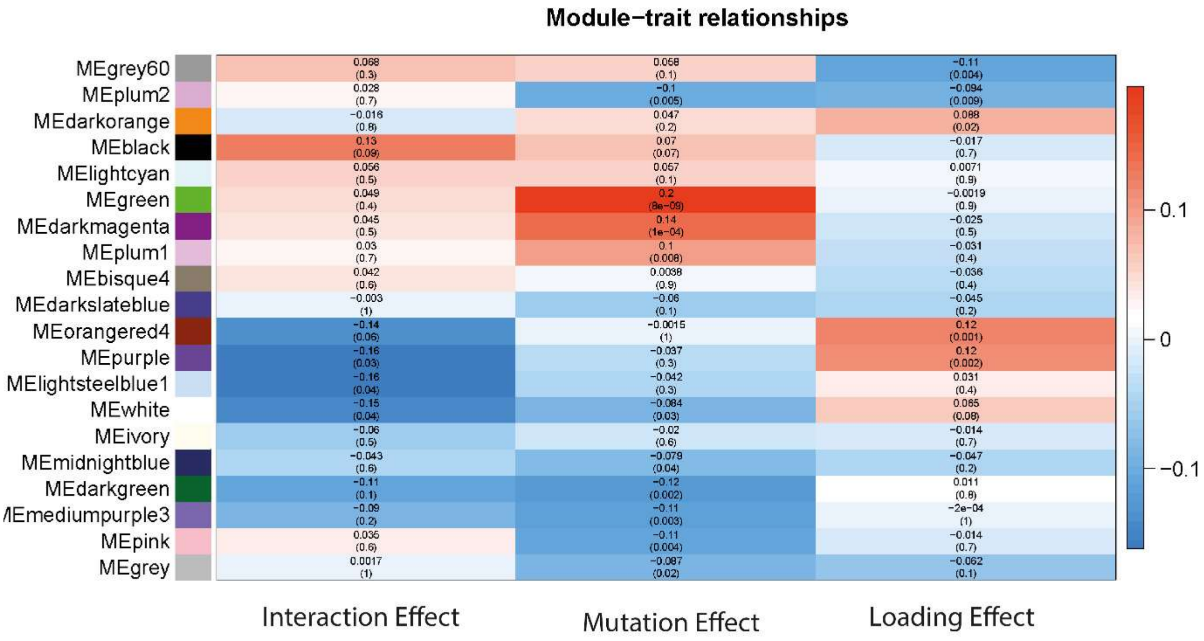


**Supplementary Figure 3.** Association of co-expression modules as determined by WGCNA with the *COL6A3* mutation, hyper-physiologic mechanical loading conditions, and the interaction between the *COL6A3* mutation and hyper-physiologic mechanical loading conditions. Data is noted as beta (p-value)

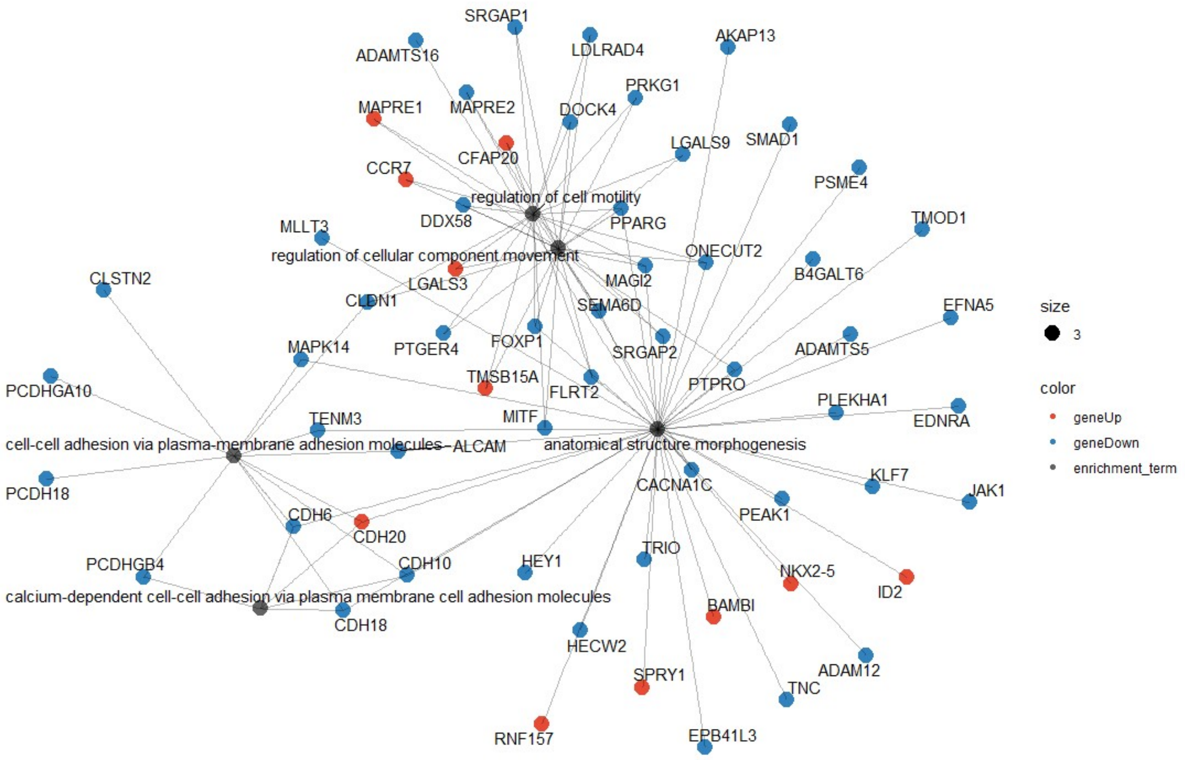


**Supplementary Figure 4.** Enrichment network of the darkgreen module associated with the *COL6A3* mutation.

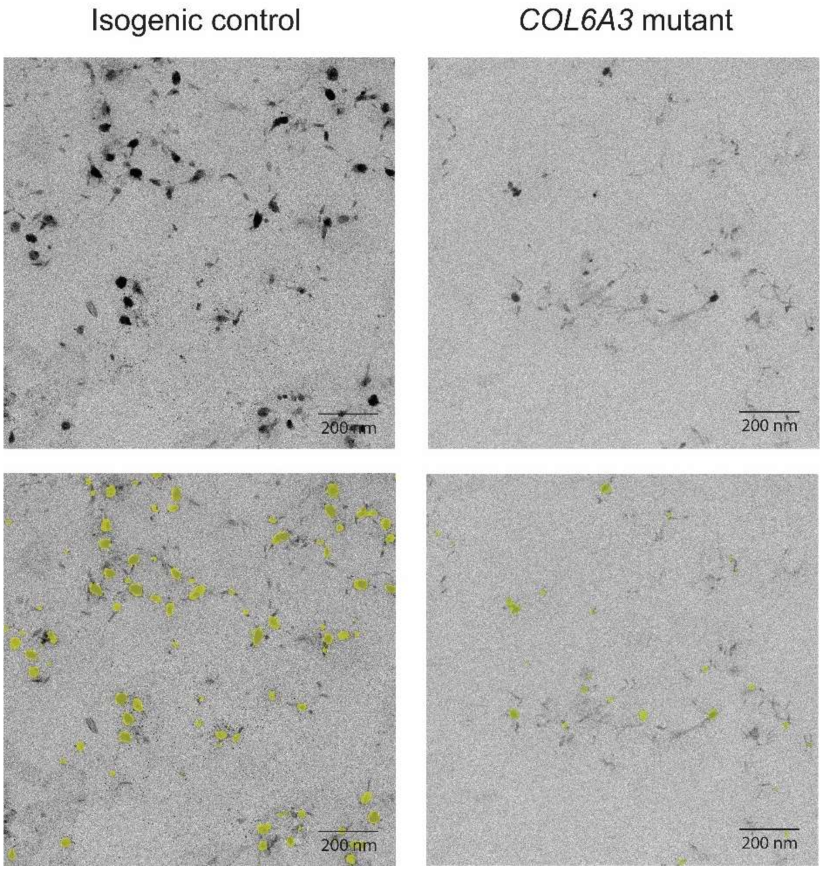


**Supplementary Figure 5.** Prediction of GAGs in TEM data. Yellow denotes the predicted annotation using a machine learning algorithm.

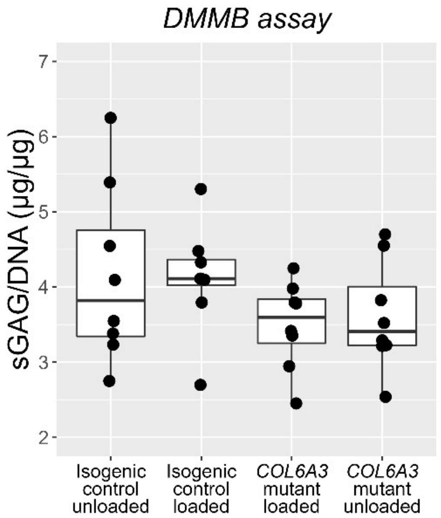


**Supplementary Figure 6. No significant effect of hyper-physiological loading conditions on loss of sGAGs.** DMMB assay of isogenic control, *COL6A3* Mutant neo-cartilage pellets, either unloaded control or exposed to hyper-physiological loading conditions.

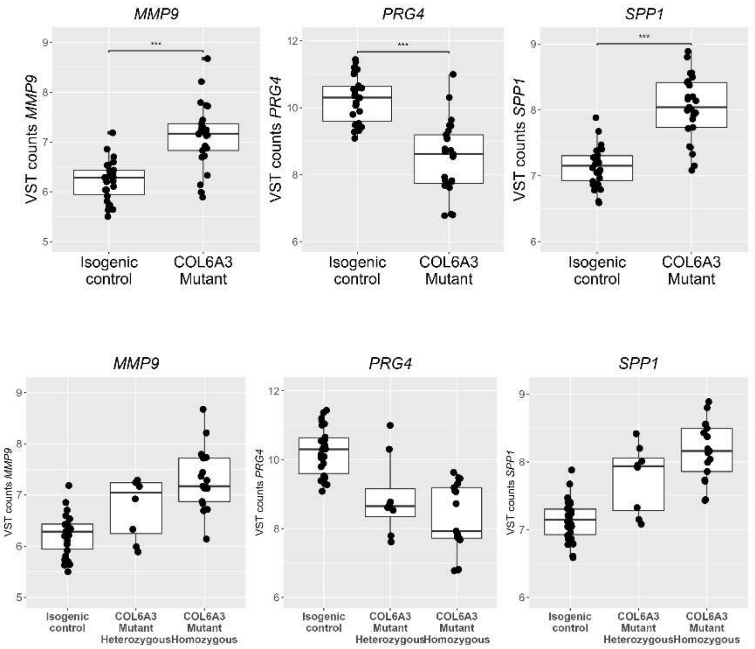


**Supplementary Figure 7.** Example of dose response effect of the hetero- and homozygous mutated *COL6A3* samples.

### Supplementary methods

*Experimental design*

The objective of the current study was to study the effects of a likely causal pathogenic mutation, thereby elucidating the effects of aberrant collagen VI functions. Exome sequencing was applied to identify a pathogenic mutation. To study the underlying effect of this mutation, the mutation was introduced in hiPSCs using CRISPR-Cas9 genome engineering, of which a neo-cartilage model was created, followed by mechanical loading and functional analysis.

*Exome sequencing*

Exome sequencing of a sibling pair with generalized OA at multiple joint sites was performed by Illumina HiSeq 2000 technology (Beijing Genome Institute). The sequences were generated as 100–base pair paired-end reads, after enrichment of 44-Mb exonic sequences by NimbleGen EZ (Roche NimbleGen). Raw imaging files were processed by Illumina base-calling software v1.7 with default parameters. SOAPaligner/SOAP2.21 was used to align reads to the GRCh37 reference genome at the UCSC Genome Browser website (http://genome.ucsc.edu/) (*34*).

*hiPSC line and cell culture*

An hiPSC line as described earlier was used as the unedited isogenic control (*19*). In short, RVR-iPSC line was retrovirally reprogrammed from BJ fibroblasts and characterized. The hiPSCs were maintained under standard conditions (37 °C, 5% $CO_2$) on Matrigel (Corning) coated plates and refreshed daily with TeSR-E8 medium (STEMCELL Technologies) upon reaching approximately 70% confluence.

*Genome editing of hiPSCs*

gRNA targeting *COL6A3* (5’-TCTGAAAACCTACAGATCCC -3’) were complexed with Cas9 protein at 25 C for 10-15’ and added to a RVR-hiPSC cell suspension of 500K cells in 400 µL mTeSR to generate the edited hiPSCs. 500 pmol of ssODN (5’-CTTGCCACAAATTCGAGAGCCTTGCCAGTG TTCAGTGGGGACCCCCCTCTGAGCCTCAGGCGCCATATGGCGTCCAGCACCGGGGC GTGGGATCTGTAGGTTTTCAGATAGAATTCTGGGAAGACAT-3’) harboring the risk variant was added to cell suspension. Cells were electroporated with BioRad Gene Pulser system with the following conditions: [250 V, 750 µF, ∞ Ω, 0.4 cm]. Cells were plated at low density and colonies were picked to obtain clones derived from single cells. As the risk variant created an NDEI site, clones were screened by PCR amplification of the risk allele and digestion with NDEI (New England Biolabs) in CutSmart Buffer for 1 hour at 37C. Successful editing of the targeted variant was confirmed by Sanger sequencing **(Fig S1)**.

*hiPSC differentiation to induced chondrocytes*

Two different chondrogenic constructs were used for downstream analysis; these chondrogenic pellets were directly used for further experiments, or they were dissociated using collagenase II, encapsulated in 2% w/v agarose at 30 million cells/ml, and cultured for 14 days with CD creating cylindrical shaped constructs. When hiPSCs reached 60% confluence, the culture medium was switched to mesodermal differentiation (MD) medium, composed of IMDM GlutaMAX (IMDM; Thermo Fisher Scientific) and Ham’s F12 Nutrient Mix (F12; Sigma-Aldrich) with 1% chemically defined lipid concentrate (Gibco), 1% insulin/human transferrin/selenous (ITS+; Corning), 0.5%

penicillin-streptomycin (P/S; Gibco), and 450 µM 1-thioglycerol (Sigma-Aldrich). Before induction of anterior primitive streak (day 0), hiPSCs were washed with wash medium (IMDM/F12 and 0.5% P/S) and then fed with MD medium supplemented with activin A (30 ng/ml; Stemgent), 4 µM CHIR99021 (CHIR; Stemgent), and human fibroblast growth factor (20 ng/ml; FGF-2; R&D Systems) for 24 hours. Subsequently, the cells were washed again with wash medium, and paraxial mesoderm was induced on day 1, by MD medium supplemented with 2 µM SB-505124 (Tocris), 3 µM CHIR, FGF-2 (20 ng/ml), and 4 µM dorsomorphin (Tocris) for 24 hours. Before induction of early somite (day 2), cells were washed with wash medium, and then cells were fed with MD medium supplemented with 2 µM SB-505124, 4 µM dorsomorphin, 1 µM C59 (Cellagen Technology), and 500 nM PD173074 (Tocris) for 24 hours. Subsequently, cells were washed with wash medium, and for induction of sclerotome, cells (days 3 to 5) were fed daily with MD medium supplemented with 2 µM purmorphamine (Stemgent) and 1 µM C59. To induce chondroprogenitor cells (days 6 to 14), cells were washed briefly with wash medium and fed daily with MD medium supplemented with human bone morphogenetic protein 4 (BMP-4; 20 ng/ml; Miltenyi Biotec). Three independent differentiations were done per clone.

Monolayer cultured hiCPC aggregates present at day 14 of the differentiation were washed with MD medium, dissociated with Gentle Cell dissociation medium (Stem Cell), and centrifuged for 5 min at 1200 rpm. Cell aggregates were subsequently maintained in chondrogenic differentiation (CD) medium containing Dulbecco's modified Eagle's medium/F12 (Gibco), supplemented with 1% ITS+, 55 µM 2-mercaptoethanol (Gibco), 1% non-essential amino acids (Gibco), 0.5% P/S, L-ascorbate-2-phosphate (50 µg/ml; Sigma-Aldrich), L-proline (40 µg/ml; Sigma-Aldrich), ML329 (1µM; CSNpharm), C59 (1µM; Tocris), and transforming growth factor–β3 (10 ng/ml; PeproTech) for 30 days while refreshing medium every 3 to 4 days.

*Mechanical loading*

The spherical shaped neo-cartilage constructs were mechanically loaded using a MACH-1 mechanical testing device (Biomomentum), at a rate of 5hz with 20% sinusoidal peak-to-peak strain for 10 minutes as described earlier (*21*)

*sGAG measurement*

Sulphated glycosaminoglycan (sGAG) concentrations in the neo-cartilage organoids (µg sGAG/µg DNA) was measured using the Farndale Dimethyl Methylene Blue (DMMB, Sigma) method (*36*). Chondroitin sulphate (Sigma) was used as a reference standard. Absorbance was measured at 535 and 595 using a microplate reader (Synergy HT, Biotek). Neo-cartilage sGAG concentrations were corrected for DNA content measured with the Qubit® 2.0 Fluorometer (Invitrogen™) using the dsDNA HS Assay Kit (Invitrogen™).

*Histology and immunohistochemistry*

Neo-cartilage samples were fixed in 4% formaldehyde and embedded in paraffin. Sections were stained with Alcian Blue (Sigma-Aldrich) and Nuclear Fast Red (Sigma-Aldrich). Deposition of collagen II and collagen VI in the neo-cartilage constructs was visualized immunohistochemically. For collagen VI, antigen retrieval was done by treating deparaffinized sections with proteinase K (5 µg/ml, Qiagen) and hyaluronidase (5 mg/ml, Sigma). Sections were incubated overnight with a primary antibody raised against human collagen VI (1:100, abcam), followed by incubation with a HRP conjugated secondary antibody (ImmunoLogic). Peroxidase binding for collagen VI was visualized using diaminobenzidine, and sections were counterstained with haematoxylin.

*RT-qPCR*

Per sample, two replicate neo-cartilage pellets were collected in TRIzol (Invitrogen™) and RNA was isolated using the RNeasy Mini Kit (Qiagen) according to manufacturer's protocol. DNA contamination was removed by treating the RNA with RNase-Free DNase RNA quality (A260/280: 1.7-2.0) was assessed using the Nanodrop. RNA concentrations were measured with the Qubit® 2.0 Fluorometer (Invitrogen™) using the RNA HS Assay Kit (Invitrogen™)., respectively, with an A260/280 between 1.7-2.0. RNA was reverse transcribed into cDNA using the Transcriptor First Strand cDNA Synthesis Kit (Roche). cDNA was amplified using FastStart SYBR Green Master (Roche) and mRNA expression was measured in triplicates in a MicroAmp™ Optical 384-Well Reaction Plate (ThermoFisher Scientific), using the QuantStudio™ Flex Real-Time PCR system (Applied Biosystems™), with the following cycling conditions: 10 min 95 °C; 10 sec 95 °C, 30 sec 60 °C, 20 sec 72 °C (45 cycles); 1 min 65 °C and 15 sec 95 °C . Primer efficiency was tested using a cDNA dilution series, and primers were considered efficient with an efficiency between 90% and 110%. -ΔCt expression levels were calculated using two housekeeping genes *GAPDH* and *SDHA*, with the following formula: *ΔCt=Ct (gene of interest) – Ct (average housekeeping genes)*. Both housekeeping genes were stably expressed in this model. Fold changes were calculated using the $2^{-\Delta\Delta Ct}$ method with *ΔΔCt=ΔCt (MS) – ΔCt (Control)*.

*RNAseq*

RNA from neo-cartilage constructs was extracted 12 hours post mechanical loading. RNA from spherical neo-cartilage constructs was extracted and processed using a pestle homogenizer in TRIzol reagent (Invitrogen). RNA was extracted using chloroform, followed by precipitation using ethanol, and purified with the RNeasy Mini Kit (Qiagen). Genomic DNA was removed by DNase digestion (Qiagen). Paired-end 2 × 150 base pair RNA sequencing (Illumina NEBNext Ultra II Directional RNA Library Prep, Illumina NOVAseq 6000) was performed. Strand-specific RNA-sequencing libraries were generated which yielded on average 25 million reads per sample. Data from the Illumina platform was analyzed with an in-house pipeline as previously described (*16*). The adapters were clipped using Cutadapt v1.1. RNA-seq reads were then aligned using GSNAP against GRCh38 (*37*). Read abundances per sample were estimated using HTSeq count v0.11.1 (*38*) with Ensembl gene annotation version 94. Only uniquely mapping reads were used for estimating expression. The quality of the raw reads and initial processing for RNA sequencing was checked using MulitQC v1.9 (*39*). Samples containing > 50% genes with zero values and average read count < 4 were removed from further analysis. The datasets from both neo-cartilage models were combined using surrogate variable analysis using the R package SVA version 3.42.0 (*40*). Outliers where identified (n=2) using hierarchical clustering and principal component analysis (PCA) which were removed from further analysis. In total, 52 samples were included of which; isogenic controls (free-swelling isogenic controls=13, mechanically stimulated isogenic controls=13) as well as COL6A3-mutant (free-swelling COL6A3 mutants=14 and mechanically loaded COL6A3 mutants, n=12) organoids. Differential expression analysis was performed using the R package DESeq version 1.34 (*41*). All samples were combined in a multifactorial analysis. To determine effects of the mutation and hyper-physiological mechanical loading conditions only the main effects were included into the model. To determine the interaction effect an interaction term to this model was added. P-values were corrected for their false discovery rate using the Benjami and Hochman method (*42*). WGCNA analysis was performed using the R package WGCNA version 1.71 (*20, 43*). A generalized linear model was used to determine the association between the COL6A3 mutation, hyper-physiological mechanical loading conditions, the interaction effect and the identified WGCNA co-expression networks. Over representation enrichment analyses of these co-expression networks using the KEGG, Reactome and gene ontologie biological processes databases was performed using the anRichment R package version 1.22. Protein-protein network analysis was performed using the online tool STRING version 11.0 (*44*).

*Solid-phase binding assay*

Conditioned medium of wild-type and *COL6A3* mutant organoids was collected and concentrated in preparation for the binding assay. To this end, 450 µl of medium was collected in 100 K molecular weight cutoff Pierce Protein Concentrators (Thermo Scientific) and centrifuged for 10 min at 12,000g. Subsequently, COL6 concentration was determined using the Human COL6A3 ELISA Kit (Assay Genie) according to the manufacturer's protocol.

Clear multiwell plates (R&D Systems) were coated overnight with 100 µl of purified fibronectin (10 µg/ml; Merck) in phosphate-buffered saline (PBS) at 4°C, followed by four wash steps with wash buffer (0.05% Tween 20 in PBS). Nonspecific binding was blocked for 1 hour with 3% (w/v) bovine serum albumin (BSA) in PBS. After washing with wash buffer, the plates were incubated with 100 µl of concentrated medium samples at COL6A3 concentration of 1.3 ng/ml in assay buffer (0.05% Tween 20 and 0.5% BSA in PBS) for 2 hours. Plates were then washed four times with wash buffer and incubated with rabbit anti-COL6A3 biotin-conjugated antibody (Assay Genie) at 0.2 µg/ml in assay buffer for 1 hour. Plates were washed, after which the plates were incubated with streptavidin–horseradish peroxidase (Thermo Scientific) at 0.1 µg/ml in assay buffer for 1 hour. After washing, color development was performed with 100 µl of tetramethylbenzidine substrate (Thermo Fisher Scientific) for 10 min, reaction was stopped with 100 µl of 1 M HCl, and absorbance was measured at 450 nm. Assays were performed in triplicate.

*Transmission electron microscopy*

To the neo-cartilage organoids, consisting of cells and matrix, double concentrated fixative was added to the culture medium resulting in a final concentration of 1,5% glutaraldehyde solution in 0,1M cacodylate buffer. The spheres were kept in fixative for at least an hour at room temperature. After rinsing 3 times with 0,1M cacodylate buffer the spheres were postfixed in 1% osmium tetroxide / 1,5% potassium ferricyanide / 0,1M cacodylate buffer at 4°C for an hour. After 3 times rinsing with 0,1M cacodylate buffer the spheres were divided into 4 quarters. The quarters were dehydrated in a series of ethanol (70%, 80%, 90% and 100% ), followed by an infiltration series of acetone / EPON LX112 (Ladd Research Industries) mixtures (2:1, 1:1 and 1:2), each step 30 min, and finally in pure EPON LX 112 for 1 hour. The quarters were positioned in a mold with the wide side towards the cutting surface, filled up with EPON and put in an 70 °C oven to polymerize for 48 hours.

Ultrathin sections (90 nm) were made with a Leica EM UC6 ultramicrotome and collected on 50 mesh grids. Sections were stained with 7% uranyl acetate in MilliQ for 10 minutes and lead citrate (*45*). Sections were then imaged in a Tecnai T12 twin (FEI / Thermo Fisher Scientific) with a Gatan 4kx4k OneView camera (Gatan) at 21.000x magnification (at a pixel size of 1.03 nm) for machine learning analysis using automated tiled imaging (MyTEM) and images were stitched together into virtual slides (MyStitch) (*46*).

*Image Analysis*

For image analysis, in total 18 virtual slides were used from three isogenic control and three *COL6A3* mutant neo-cartilage organoids (3 positions each). Using a custom-written user interface software (to be published) GAG structures were manually annotated in both control and mutated samples and from this a model was trained for machine learning. The structures were predicted in all 18 slides (Fig. S5). Both the surface of the GAGs and their number was extracted from the predictions.
